# Seasonal stability of SARS-CoV-2 in biological fluids

**DOI:** 10.1101/2021.04.07.438866

**Authors:** Taeyong Kwon, Natasha N. Gaudreault, Juergen A. Richt

## Abstract

Transmission of SARS-CoV-2 occurs by close contact with infected persons through droplets, the inhalation of infectious aerosols and the exposure to contaminated surface. Previously, we determined the virus stability on different types of surfaces under indoor and seasonal climatic conditions. SARS-CoV-2 survived the longest on surfaces under winter conditions, followed by spring/fall and summer conditions, suggesting the seasonal pattern of stability on surfaces. However, under natural conditions, the virus is secreted in various biological fluids from infected humans. In this respect, it remains unclear how long the virus survives in various types of biological fluids. This study explored the SARS-CoV-2 stability in human biological fluids under different environmental conditions and estimated the half-life. The virus was stable for up to 21 days in nasal mucus, sputum, saliva, tear, urine, blood, and semen; it remained infectious significantly longer under winter and spring/fall conditions than under summer conditions. In contrast, the virus was only stable up to 24 hours in feces and breast milk. These findings demonstrate the potential risk of infectious biological fluids in SARS-CoV-2 transmission and have implications for its seasonality.

## 1. Introduction

Since its first emergence in China, SARS-CoV-2 has rapidly spread worldwide and posed a tremendous threat to global public health and economy. The efficient transmission of SARS-CoV-2 is primarily mediated through the close contact with infected individuals who shed respiratory droplets when exhaling, talking, sneezing and coughing [1]. The expelled droplets span a wide size spectrum, but the majority of them are generally considered > 5 μm in diameter. In the contrast, in airborne transmission, smaller droplets, size ≤ 5 μm, evaporate quickly to generate infectious aerosols that can remain suspended in air for several hours and travel over a long distance [2]. Third route of transmission is the exposure to the infectious virus on surfaces. Large droplets settle down to the ground or surfaces within a limited distance and contaminate the environment with the virus. The infectious virus can survive for several days on various types of surfaces and the half-life is the longest under winter conditions, followed by spring/fall and summer conditions, suggesting a seasonal pattern of virus survival on contaminated surfaces [3, 4]. Oro-fecal/naso-fecal, bloodborne, mother-to-child, and sexual transmissions have also been proposed, but their roles still remain unclear.

Scientists have investigated the presence of SARS-CoV-2 in biological fluids since it serves as a potential source of infection and clinical specimens for diagnostics. Nasal mucus, sputum and saliva are major components to produce the respiratory droplets by which the transmission of SARS-CoV-2 predominantly occurs. Ocular changes and conjunctivitis are one of the clinical manifestations of SARS-CoV-2 infection in humans, and the virus can be detected and isolated in tears and ocular swabs [5]. In rare cases, infectious virus was isolated in urine [6] and feces [7–9] of COVID-19 patients and these findings suggest a possible route of oro-fecal/naso-fecal transmission of SARS-CoV-2, even transmission through the wastewater system. Several studies reported the detection of SARS-CoV-2 RNA in blood [10, 11], breast milk [12, 13] and semen [14] which might suggest bloodborne, mother-to-child and sexual transmission, respectively. Therefore, we extensively evaluated the virus stability in liquid and surface settings of human biological fluids under indoor and different seasonal conditions.

## 2. Results

We estimated SARS-CoV-2 half-life (t_1/2_) values in human nasal mucus, sputum, saliva, tears, blood and semen under both liquid and a non-porous surface setting, as well as human urine in liquid only. Under indoor conditions (21 °C/60% relative humidity [RH]), t_1/2_ values ranged from 5.23 to 16.74 hours in liquid and from 6.77 to 16.57 hours on a steel surface (Table 1). We found t_1/2_ values ranging from 2.3–12.57 and 2.58–10.75 hours in liquid and surface settings, respectively, under summer conditions (25 °C/70% RH). The incubation under spring/fall conditions (13 °C/66% RH) resulted in slower virus decay and longer t_1/2_ values resulting in 15.98–54.34 hours in liquid and 18.15–48.4 hours for the surface setting. The longest virus survival was observed under winter conditions (5 °C/75% RH), where t_1/2_ values were 33.37–121.83 hours in liquid and 38.55–235.18 hours for the steel surface setting. Statistical analysis on liquid and surface settings showed that the t_1/2_ values under summer conditions were significantly different from either the spring/fall or winter conditions (adjusted *p* < 0.0001) for human nasal mucus, sputum, saliva, tears, blood and semen, and for human urine in liquid (Figure 1). In addition, we found significant differences between spring/fall and winter conditions for human saliva (adjusted *p* = 0.0331) and tears (adjusted *p* = 0.0029) in liquid, and for tears (adjusted *p* = 0.0146) and blood (adjusted *p* = 0.0202) under the surface setting. SARS-CoV-2 was more stable in the liquid setting than surface setting for nasal mucus under summer conditions, and for tears under indoor and summer conditions (Table 1). In contrast, the t_1/2_ values were significantly higher on the surface than liquid for sputum under indoor, spring/fall and winter, for saliva under indoor and spring/fall, for blood under winter, and for semen under summer, spring/fall and winter conditions.

**Table 1.** Half-life of SARS-CoV-2 in human biological fluids under indoor, summer, spring/fall and winter conditions.

| Environmental condition | 21 °C/60% RH indoor |  | 25 °C/70% RH summer |  | 13 °C/66% RH spring/fall |  | 5 °C/75% RH winter |  |
| --- | --- | --- | --- | --- | --- | --- | --- | --- |
| Setting | Liquid | Surface | Liquid | Surface | Liquid | Surface | Liquid | Surface |
| Nasal mucus | 5.23<br>(4.03, 7.47)<sup>1</sup> | 6.77<br>(4.57, 13.01) | 4.59<sup>2</sup><br>(3.66, 6.17) | 2.58<sup>2</sup><br>(2.02, 3.59) | 21.74<br>(15.61, 35.76) | 18.15<br>(14.66, 23.83) | 53.94<br>(44.82, 67.71) | 38.55<br>(30.42, 52.62) |
| Sputum | 8.69<sup>2</sup><br>(7.11, 11.17) | 14.9<sup>2</sup><br>(10.72, 24.37) | 3.68<br>(2.59, 6.37) | 5.55<br>(4.42, 7.45) | 21.94<sup>2</sup><br>(16.05, 34.66) | 37.03<sup>2</sup><br>(27.74, 55.67) | 33.37<sup>2</sup><br>(22.75, 62.56) | 76.4<sup>2</sup><br>(60.48, 103.7) |
| Saliva | 7.89<sup>2</sup><br>(6.38, 10.37) | 12.69<sup>2</sup><br>(10.36, 16.38) | 6.98<br>(5.19, 10.65) | 6.44<br>(4.87, 9.5) | 15.98<sup>2</sup><br>(12.51, 22.12) | 34.17<sup>2</sup><br>(27.05, 46.28) | 55.16<br>(44.28, 73.15) | 69.25<br>(61.3, 79.57) |
| Tears | 15.1<sup>2</sup><br>(12.29, 19.56) | 8.3<sup>2</sup><br>(7.09, 10) | 11.06<sup>2</sup><br>(8.91, 14.58) | 3.53<sup>2</sup><br>(2.93, 4.45) | 29.34<br>(25.75, 34.07) | 24.22<br>(20.14, 30.38) | 121.83<br>(91.19, 183.44) | 106.82<br>(84.44, 145.43) |
| Urine | 11.41<br>(9.69, 13.88) | N/A<sup>3</sup> | 7.89<br>(6.74, 9.5) | N/A<sup>3</sup> | 54.34<br>(36.76, 103.8) | N/A<sup>3</sup> | 57.73<br>(49.06, 70.15) | N/A<sup>3</sup> |
| Blood | 16.74<br>(11.64, 29.83) | 16.57<br>(13.1, 22.53) | 12.57<br>(9.58, 18.3) | 10.75<br>(9.16, 13) | 39.25<br>(28.03, 65.26) | 48.4<br>(35.79, 74.6) | 102.04<sup>2</sup><br>(71.93, 175.63) | 235.18<sup>2</sup><br>(144.52, 631.75) |
| Semen | 7.48<br>(5.76, 10.65) | 9.51<br>(7.46, 13.1) | 2.3<sup>2</sup><br>(1.75, 3.33) | 5.9<sup>2</sup><br>(4.75, 7.78) | 17.69<sup>2</sup><br>(13.96, 24.14) | 41.24<sup>2</sup><br>(29.31, 69.31) | 57.81<sup>2</sup><br>(47.77, 73.23) | 91.64<sup>2</sup><br>(75.24, 117.22) |
| Positive control | 15.85<sup>2</sup><br>(11.81, 24.12) | 7.88<sup>2</sup><br>(6.48, 10.05) | 7.48<sup>2</sup><br>(6.7, 8.46) | 2.57<sup>2</sup><br>(2.21, 3.08) | 48.95<sup>2</sup><br>(35.25, 80) | 21.5<sup>2</sup><br>(17.25, 28.56) | 176.66<sup>2</sup><br>(109.78, 451.39) | 79.64<sup>2</sup><br>(63.6, 106.48) |

<sup>1</sup> Half-life (95% confidence interval) in hours
<sup>2</sup> Significant difference ( $p < 0.05$ ) between liquid and surface settings under a respective condition
<sup>3</sup> N/A: Not available

**Figure 1.**
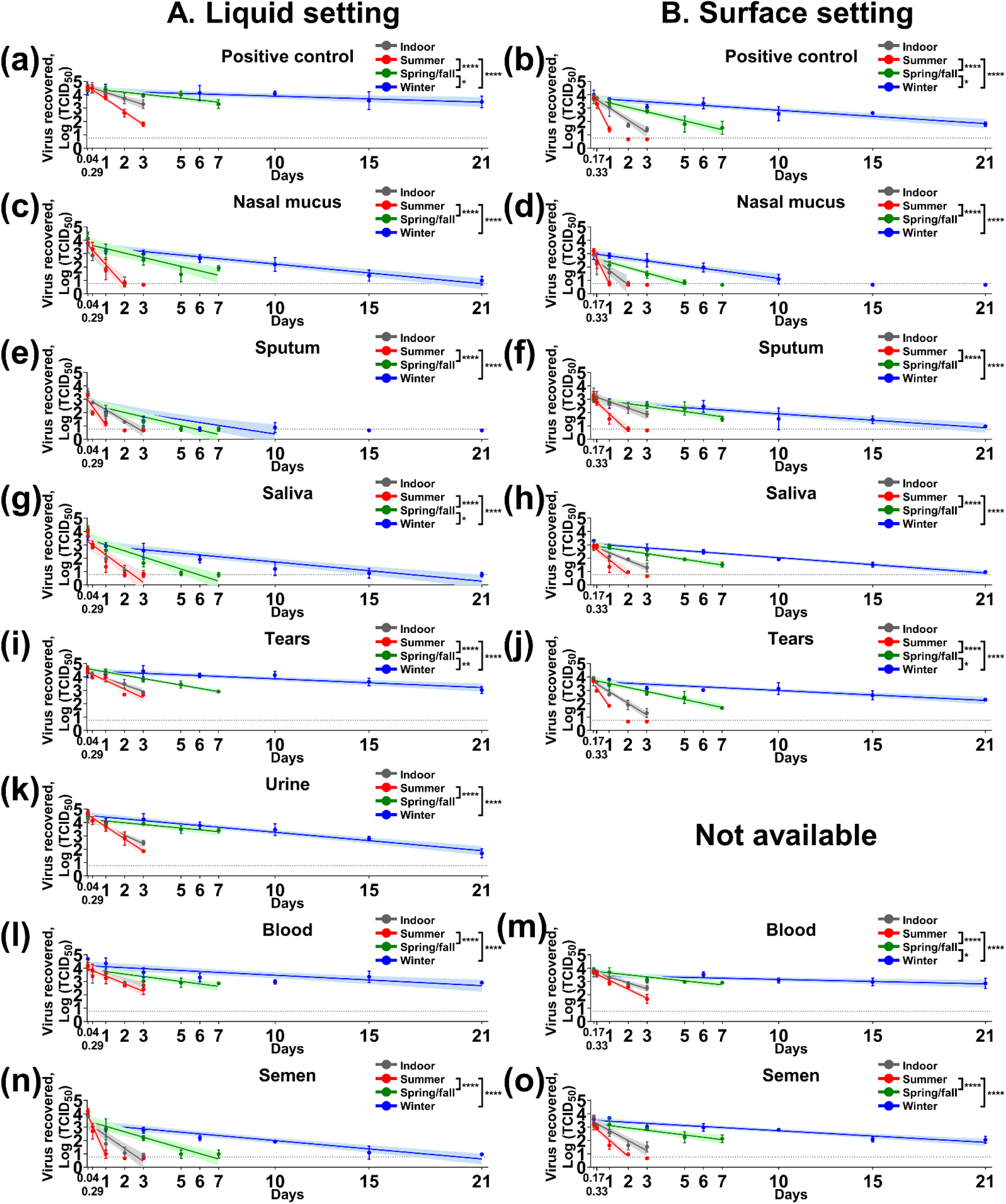
SARS-CoV-2 stability in medium, human nasal mucus, sputum, saliva, tears, urine, blood, and semen under indoor and three seasonal conditions. The mixture of the virus (5×10^4^ 50% tissue culture infectious dose [TCID_50_]) and each biological fluid was placed in sealed tubes for (A) liquid setting (a, c, e, g, i, k, l and n) or (B) stainless steel for the surface setting (b, d, f, h, j, m and o). The mixture was dried for 4 hours inside a biosafety cabinet for the surface setting. The tubes and stainless steel were incubated under indoor (grey), summer (red), spring/fall (green) and winter (blue). At each time point, infectious virus was recovered in 2 mL medium, filtered through a 0.45 μm syringe filter and titrated on Vero E6 cells. Virus titers were log-transformed and a simple linear regression model was determined in Prism 9, Graphpad. Virus titers are expressed as mean log TCID_50_ ± standard deviation at each time point; the solid lines and theirs shade areas represent a best-fit line and 95% confidence interval of linear regression model under each condition. The dash line represents the limit of detection, 10^0.767^ TCID_50_, for the virus isolation assay. Statistical analysis using ANOVA and subsequent Tukey’s adjustment was performed to determine seasonal difference of t_1/2_ half-life of SARS-CoV-2 in summer, spring/fall and winter conditions. Significance was marked * (adjusted *p* < 0.05), ** (adjusted *p* < 0.01), *** (adjusted *p* < 0.001) and **** (adjusted *p* < 0.0001). On x-axis, 0.04, 0.17, 0.29 and 0.33 days are equal to 1, 4, 7, and 8 hours, respectively.

However, the virus was unstable under both liquid and surface settings in human feces, fecal suspension, and breast milk, as well as in urine under the surface setting. In human fecal suspension, low virus titers were found at 1 hour post-contamination (hpc) in liquid under indoor (0.978 ± 0.317 log 50% tissue culture infectious dose [TCID_50_]) and summer conditions (0.767 ±0.174 log TCID_50_). No infectious virus was recovered from human feces for either setting, or from fecal suspension and urine on the steel surface. In both settings of human breast milk, the infectious virus was detected up to 1 day post-contamination (dpc), in which virus titers ranged from 0.767 ± 0.174 to 1.812 ± 1.151 log TCID_50_ (Figure 2).

**Figure 2.**
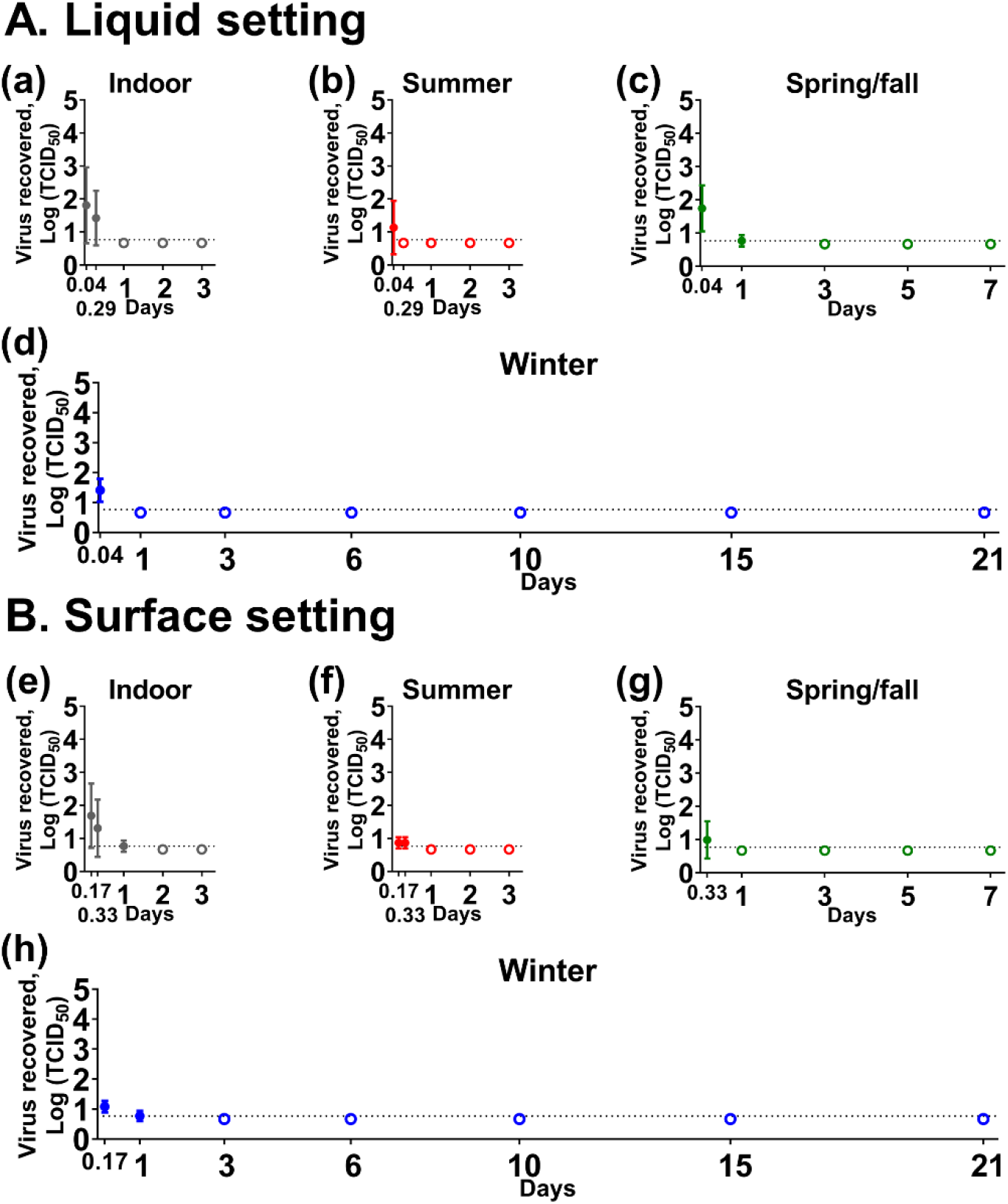
SARS-CoV-2 stability in human breast milk. A mixture of the virus (5×10^4^ 50% tissue culture infectious dose [TCID_50_]) and breast milk was placed in sealed tubes for (A) the liquid setting (a–d) or on (B) stainless steel for surface setting (e–h) and incubated under indoor (a and e), summer (b and f), spring/fall (c and g) and winter (d and h) conditions. Virus titer was expressed as mean log TCID_50_ ± standard deviation at each time point. The dash line represents a limit of detection, 10^0.767^ TCID_50_, for the virus isolation assay. Empty dots represent negative results tested in triplicate. On x-axis, 0.04, 0.17, 0.29 and 0.33 days are equal to 1, 4, 7, and 8 hours, respectively.

## 3. Discussion

Extended stability and seasonality are one of the intrinsic characteristics of SARS-CoV-2, which could potentially contribute to spread the virus through contaminated surfaces and accelerate the transmission in cold weather. Earlier studies have investigated the virus stability by contaminating various surfaces with cell culture-derived virus inoculum [3, 4, 15]. The virus remained infectious on various surfaces for several days under indoor conditions and its survival was significantly longer under winter climatic conditions than summer climatic conditions [3]. Some studies added bovine serum albumin (BSA) or a mixture of BSA, mucin and tryptone into virus inoculum used for surface contamination, in order to mimic the proteinaceous and organic content of body fluids; this resulted in an extended infectivity period on surfaces of up to 28 days after contamination at 20 °C [16, 17]. However, the limitations of this approach are that the content of organic matter is dependent on the type of biological fluids and that fluids contain substances which could be beneficial or harmful for virus survival. These drawbacks led to new approaches to test SARS-CoV-2 stability in biological fluids, such as human nasal mucus, sputum, fecal suspension and urine [18, 19].

In the present study, we determined the potential risk of biological fluids to spread SARS-CoV-2 by testing its stability in nine human biological fluids from which the virus has been isolated (nasal mucus, sputum, saliva, tears, feces and urine) or detected (breast milk, blood and semen). Generally, SARS-CoV-2 is excreted in biological fluids from infected patients. It can contaminate surfaces and eventually dry on surfaces as the result of water evaporation. The duration of evaporation is dependent on various factors, such as the volume of water and surrounding environmental conditions. Therefore, we evaluated virus stability in various biological fluids using (1) a liquid setting in which the mixture was placed in a sealed tube to prevent evaporation and (2) a surface setting in which the mixture was completely dried on the stainless steel surface. Both beneficial and harmful substances concentrate when dried on the surface, and there is little or no activity of microorganisms, broadly active viral inhibitors, or enzymes due to the lack of water in the surface setting. In the present study, the virus-spiked biologicals in sealed tubes and on stainless steel surfaces were incubated under indoor and three seasonal conditions which represent U.S. Midwest climate conditions.

Nasal mucus, sputum and saliva are major components of droplets which play a primary role in transmission of SARS-CoV-2. Infectious droplets which are generated by infected individuals can reach the mouth and nose of a susceptible individual within a close distance. However, the droplet can also contaminate surfaces and remain infectious for a certain period of time. The potential risk of fomites in SARS-CoV-2 transmission has motivated the investigation of virus stability in biological fluids. A previous study showed that SARS-CoV-2 survived in nasal mucus and sputum for 24 hours under both indoor and winter conditions [18]. However, our data demonstrated that the virus was able to remain infectious for at least 2 days, 2 days, 7 days and 21 days in nasal mucus and 3 days, 2 days, 7 days and 21 days in sputum under indoor, summer, spring/fall and winter conditions, respectively. Also, SARS-CoV-2 survived in saliva for a similar time period. The disparity in the viral stability might be explained by the different sources of biological fluids, different inoculum titers and different experimental settings. In the present study, we also demonstrated the effect of temperature and relative humidity on virus survival and found that virus stability was significantly extended under spring/fall and winter, but not summer conditions. These results imply the seasonal pattern of SARS-CoV-2 stability in nasal mucus, sputum and saliva. After the first wave of COVID-19 pandemic took place in spring of 2020, we faced a small rise in the number of new coronavirus infection in June across the planet, which might be associated with a relaxation of lockdowns and an alleviation of the public’s awareness. However, in the fall and winter of 2020/2021, there has been a drastic re-surgence in new daily COVID-19 cases in the Northern Hemisphere. Along with socioeconomical factors, such as COVID-fatigue and more indoor gatherings during colder temperatures, and the emergence of new variants, the seasonality of SARS-CoV-2 stability in biological fluids might be a factor associated with enhanced transmission which started in the late fall of 2020.

Despite the tissue and cellular tropism of SARS-CoV-2 in conjunctiva *ex vivo* [20], the role of tears in transmission has been underestimated due to fewer clinical manifestations and less virus detection/isolation in conjunctival excretion and tears [21, 22]. It is shown that the eye has a mucosal defense system where lysozyme, lactoferrin, lipocalin and secretory immunoglobulin A are active components in tears to exert their antimicrobial activity [23]. Lactoferrin has been regarded as a potential inhibitor to SARS-CoV-2 infection because it inhibits SARS-CoV-1 entry into cells by binding to cellular heparin sulfate [24] and SARS-CoV-2 also utilizes heparin sulfate to facilitate the attachment of the spike protein to the angiotensin-converting enzyme 2 receptor [25]. Interestingly, our results show that tears can serve as a viable matrix for virus stability, especially in the liquid setting. It is hypothesized that once the virus replicates in conjunctiva tissue and is excreted in tears, it could survive in the environment for an extended time period.

SARS-CoV-2 shedding in feces and urine has been of concern because of the potential transmission via oro-fecal/naso-fecal as well as through wastewater. To date, a few studies found viable SARS-CoV-2 in urine and stool specimens of COVID-10 patients [6–9]. Our results show that low amount of virus was detected in human fecal suspension at 1 hpc and no infectious virus in human feces throughout the study, suggesting the virus is not stable in human feces. It is plausible that infectious virus might survive by suspending a fecal swab or stool specimens in a virus transport media and inoculating it onto cells immediately. In contrast, the virus was stable in urine and survived longer at lower temperatures. These results are comparable to a previous study, in which the virus remained infectious for several hours in a fecal suspension and for three to four days in urine at room temperature [19]. In addition, no viable virus was isolated from the steel surfaces contaminated with feces, fecal suspension and urine at different time points post-contamination in this study. This could be explained by concentrated virucidal substances destroying the viable virus during the process of drying. Further analysis is needed to evaluate the potential risk of transmission through wastewater or byproduct from sewage, such as biosolids, since most human feces and urine enter the wastewater system to be processed in a sewage treatment plant [26].

SARS-CoV-2 RNAemia has been frequently found in patients suffering a severe course of infection, i.e., the presence of RNA in blood could be a potential factor in predicting critical outcome [27]. In patients with RNAemia, severe illness is considered the consequence of a dysregulation of the host immune response which is triggered by increased viral load in the serum [28]. SARS-CoV-2 in the blood has been of great interest in terms of virus transmission as the blood could serve as the source of infection through blood transfusions. However, so far, no evidence supports the transmission of SARS-CoV-2 via blood transfusion [29]. Our study, however, raises the concern that blood could be also involved in indirect transmission of SARS-CoV-2. Both, liquid and dried blood provided the most beneficial condition to stabilize SARS-CoV-2 infectivity. The high concentration of proteinaceous substances, such as bovine albumin serum, may have a positive effect on virus stability as observed in a previous study [16].

Similarly, semen has a high concentration of protein contents derived from prostatic origin and seminal fluids [30]. This physicochemical property of semen might explain the exceptional virus stability in semen observed on surfaces, especially under spring/fall and winter conditions. In the contrast, a low virus half-life was found for semen in liquid under summer conditions. Previously, it was shown that human immunodeficiency virus is inactivated in semen by at least two different mechanisms: the antiviral activity of H_2_O_2_ which is generated from the oxidation of seminal plasma by diamine oxidase [31]; and the intrinsic virucidal activity of cationic polypeptides [32]. These enzymatic and inhibitory activities in semen might account for the faster inactivation of SARS-CoV-2 in the liquid setting, especially under summer conditions.

Vertical transmission of SARS-CoV-2 via breast milk is still discussed controversially [33, 34]. Our study showed that infectious virus could be isolated from breast milk up to 1 dpc in both liquid and surface settings regardless of environmental conditions, implying the low risk of breast milk in fomite transmission. It has been of interest whether breast milk can act as a vehicle to spread viral diseases. For this purpose, stability studies of many viruses, including flavi-, Zika, and hepatitis C viruses, in breast milk were performed. Breast milk has been shown to reduce Zika virus infectivity in a time-dependent manner, with antiviral activity increasing with incubation time [35]. The antiviral activity of breast milk was closely related to the release of free fatty acids by an endogenous lipase, which resulted in the destruction of the envelope layer of hepatitis C virus and finally the disruption of viral integrity and infectivity [36]. Thus, a similar mechanism resulting in disruption of the viral envelope might explain the instability of SARS-CoV-2 in breast milk.

There are several main limitations of this study. First, no fresh biological fluid was tested in this study. Most biological fluids were collected and frozen at −20 °C. Only blood was kept at 4 °C before use. The pre-storage of biological fluid under frozen conditions before usage in the assay might have an effect on antiviral factors, which was reported for breast milk [35]. Secondly, the duration of the experiment was relatively short (maximum 21 days); we could not observe the complete disappearance of infectious virus in some of the experimental conditions. Nonetheless, we successfully calculated the biological half-life of SARS-CoV-2 in biological fluids and further determined the seasonality. Moreover, biological fluids used in this study were collected from healthy individuals. The physicochemical property of biological fluids could be different in sick individuals, which might result in different virus stability patterns. In addition, the virus stability in biological fluids has been shown to vary greatly between donors [35]; therefore, further in-depth analyses are needed to establish more precise calculations of virus half-life in biological fluids from different donors (healthy and sick) under various conditions. Lastly, we used USA-WA1 isolate which was isolated from the first U.S. patient in January 2020. However, current circulating strains have accumulated point mutations throughout the genome over time, which might have impact on the virus stability.

In summary, this study presents the SARS-CoV-2 stability in human biological fluids under indoor and three seasonal conditions. The virus was stable in nasal mucus, sputum, saliva, tears, urine, blood, and semen with t_1/2_ values of 5.23–16.74, 2.3–12.57, 15.98–54.34 and 33.37–235.18 hours under indoor, summer, spring/fall and winter conditions, respectively. The virus survival was significantly longer under either spring/fall or winter than summer conditions, suggesting seasonal effects on the stability of SARS-CoV-2 in biological fluids. Interestingly, we found that SARS-CoV-2 is unstable in human feces and human breast milk with infectious virus detected only up to 24 hours post-contamination. Our findings provide new insights into the potential role of biological fluids in SARS-CoV-2 transmission and contribute to the development of public health strategies to mitigate the risk of fomites in SARS-CoV-2 transmission.

## 4. Materials and Methods

Substrates used in this study were nine human biological fluids; nasal mucus, sputum, saliva, tears, feces, urine, breast milk, blood, and semen (Lee Biosolutions, Inc., Maryland Heights, MO, USA). Biological fluids were stored at −20 °C with the exception of blood which was kept at 4 °C before use. In addition, 10% (w/v) of fecal suspension was prepared in phosphate-buffered saline. Each test sample consisted of 5 μL of 5×10^4^ TCID_50_ SARS-CoV-2/USA-WA1/2020 virus and 45 μL of each respective substrate. The mixture was placed in a sealed tube to test stability in liquid and onto stainless steel (1/2 inch in diameter and 16 ga thickness, Metal Remnant Inc., Salt Lake City, UT, USA) in a 12-well plate for testing stability on a surface. For nasal mucus, sputum and feces which were not pipetted, 0.10 g to 0.15 g of each substrate was placed in the tube or on the stainless steel, spiked with the same amount of virus and mixed gently with a pipette tip. As a positive control for each setting, we used the same volume of Dulbecco’s Modified Eagle’s Medium (DMEM) supplemented with 5% fetal bovine serum (FBS) instead of the substrate. The virus-spiked substrates on stainless steel were air-dried for 4 hours inside a biosafety cabinet to recapitulate dried infectious biological material on a non-porous surface. The liquid virus in the sealed tube and the dried virus on the stainless steel were incubated in an environmental chamber (Temperature test chamber, Nor-Lake Scientific, Hudson, WI, USA) under four different environmental conditions; 21 °C/60% RH, 25 °C/70% RH, 13 °C/66% RH and 5 °C/75% RH, which simulates indoor, summer, spring/fall and winter conditions, respectively [3]. The infectious virus was recovered at the respective time point in 2 mL DMEM supplemented with 5% FBS, filtered through 0.45 μm syringe filter (TPP, Trasadingen, Switzerland) and titrated on Vero E6 cells (Table 2). The assay were performed in triplicate. Log-transformed virus titers in nasal mucus, sputum, saliva, tears, urine, blood, semen, and positive control were incorporate to estimate a simple linear regression in Prism 9 (GraphPad, San Diego, CA, USA). Half-life (t_1/2_) was calculated as a −log_10_(2)/slope. To determine the seasonal pattern of stability, one-way analysis of variance (ANOVA) was used to compare the slopes of linear regression under summer, spring/fall and winter conditions according to the software’s instruction. Multiple pairwise comparisons using Tukey’s adjustments were subsequently performed to identify significant difference between two conditions. Significant difference between liquid and surface settings was tested using default analysis which is compatible to analysis of covariance in GraphPad Prism 9.

**Table 2.** A series of time points at which the infectious virus was collected and titrated on Vero E6 cells after incubation under indoor or different seasonal condition.

| Environmental condition | 21 °C/60% RH/indoor, 25 °C/70% RH/summer | 13 °C/66% RH/spring/fall | 5 °C/75% RH/winter |
| --- | --- | --- | --- |
| Liquid setting | 1 hour post-contamination (hpc), 7 hpc, 1 day post-contamination (dpc), 2 dpc and 3 dpc | 1 hpc, 1 dpc, 3 dpc, 5 dpc and 7 dpc | 1 hpc, 1 dpc, 3 dpc, 6 dpc, 10 dpc, 15 dpc and 21 dpc |
| Surface setting | 4 hpc, 8 hpc, 1 dpc, 2 dpc and 3 dpc | 4 hpc, 1 dpc, 3 dpc, 5 dpc and 7 dpc | 4 hpc, 1 dpc, 3 dpc, 6 dpc, 10 dpc, 15 dpc and 21 dpc |

## Author Contributions

Conceptualization, T.K., N.N.G. and J.A.R.; methodology, T.K., N.N.G. and J.A.R.; formal analysis, T.K.; investigation, T.K.; writing—original draft preparation, T.K.; writing—review and editing, T.K., N.N.G. and J.A.R.; supervision, J.A.R.; funding acquisition, J.A.R. All authors have read and agreed to the published version of the manuscript.

## Funding

Funding for this study was provided through grants from the National Bio and Agro-Defense Facility (NBAF) Transition Fund from the State of Kansas, the MCB Core of the National Institute of General Medical Sciences (NIGMS) of the National Institutes of Health under award number P20GM130448, and the NIAID Centers of Excellence for Influenza Research and Surveillance under contract number HHSN 272201400006C. This research has also been supported by German Federal Ministry of Health (BMG) COVID-19 Research and development funding to WHO.

## Institutional Review Board Statement

Experiments were approved and performed under the Kansas State University (KSU) Institutional Biosafety Committee (IBC, Protocol #: 1452).

## Informed Consent Statement

Not applicable.

## Acknowledgments

The following reagent was obtained through BEI Resources, National Institute of Allergy and Infectious Diseases (NIAID), National Institutes of Health (NIH): SARS-CoV-2 Virus strain USA-WA1/2020 (catalogue # NR-52281).

## Conflicts of Interest

All authors declare no conflict of interest.

## Disclaimer

The authors alone are responsible for the views expressed in this publication and they do not necessarily represent the views, decisions or policies of WHO.

## Notes

### Competing Interest Statement

The authors have declared no competing interest.

